# Early stage of life is characterized by increased excitability of the auditory cortex in both humans and rats

**DOI:** 10.1101/2023.12.08.570872

**Authors:** Krista Lehtomäki, Jari Keinänen, Lauri Parkkonen, Riaz Uddin Mondal, Markku Penttonen, Tiina Parviainen

## Abstract

Auditory evoked response (ER) undergoes notable changes during childhood and likely reflects changes in synaptic signaling in the auditory cortex. Establishing the interspecies generalizability of electrophysiological indicators of cortical maturation would offer a means to better understand the neurodevelopment of the human brain. We measured cortical ERs to simple auditory probes in three age groups of human subjects and juvenile rats. These two species exhibited a remarkably similar long-latency (150–350 ms poststimulus) response in the auditory cortex, specifically pronounced in the younger individuals in both species. An age-dependent pattern of activity was evident at the level of single trials, and the late response showed stronger trial-by-trial stability in children than the early, adult-like 100-ms response, especially in humans. This robust development-related pattern of sensory cortex excitability is likely to represent a distinct synaptic event and may be a marker of the maturational stage, especially in GABAergic cortical circuits.

---

Cortical development proceeds as an interplay between genetically driven maturational processes and environmental influence. The nervous system is modified remarkably during sensitive periods in early life (Johnson 2001; Johnson 2005). This capacity of cortical plasticity underlies the adaptation of the brain’s wiring in response to the unique growth environment, thus providing the basis for a mature brain network and behavior. Despite its essential role in structuring the cerebral cortex, surprisingly little is known about the developmental trajectory of brain–environment interactions or, more specifically, the excitability of cortical circuitry to external stimulation, especially in humans.

Research of auditory perception on humans and nonhuman species that utilize hearing in social communication provides insights into our understanding of the development of behaviorally meaningful neural processes. In humans, auditory cortex electrophysiology shows notable changes across development in noninvasively recorded electrophysiological signals from outside the skull (Albrecht et al. 2000). In animal models, it has been possible to scrutinize the neurophysiological changes at the level of cortical circuitry across age using invasive recordings (Takesian and Hensch 2013), but no information exists on the extracortical response pattern in the juvenile stage. We aimed to determine whether the developmentally specific features of auditory responses represent general characteristics of the developing circuitry by implementing the same paradigm across maturing humans and rats. If comparable response characteristics in juvenile brains can be verified with parallel measurement protocols across mammalian species, there will be considerable prospects for a better understanding of cortical development and plasticity in humans.

The auditory system converts the air pressure changes received by the ear into patterns of neural activity that ultimately support all auditory-based behavior in humans and other species. The sequence of deflections in the auditory evoked response, measured by time-sensitive neuroimaging methods such as electroencephalography (EEG) and magnetoencephalography (MEG), reflects this multiphase processing in the brain (Picton et al. 1974; Hari 1990). The neural computation underlying these externally measured responses can be quantified and extrapolated by the polarity, i.e., the direction of the underlying neural current, spatial distribution, amplitude, and timing of individual deflections. Automatic (obligatory) responses to auditory input arise ∼0–200 ms after the onset of a sound and, similarly to early somatosensory responses (Stephani et al. 2021), can be seen to reflect mostly physical characteristics of the stimulus and the excitability level of the cortex (Bruyns-Haylett et al. 2017). Thus, simple auditory stimulation can be used to probe the reactivity properties, or excitability, of the neural circuitry across the auditory pathway. Indeed, the functional properties of cortical responses to simple stimulation can be thought of as reflecting the structural characteristics of the processing pathway, as, for example, the latencies and spatial distribution are directly defined by the micro- and macro-level connectivity pattern of the cortex.

The early auditory brain stem responses at 1–10 ms poststimulus (Johnson et al. 2005) are followed by middle-latency responses at around 10–60 ms (Özdamar and Kraus 1983; McGee and Kraus 1996), which are generated by various thalamocortical brain structures (Kraus and McGee 1992). Characteristic of the late auditory responses starting at around 50 ms onward is that while they signify the cortical processing sequence engaged in processing any auditory input, they also associate with perceptual and cognitive operations (Picton 2011). Indeed, at these latencies, source modeling of the underlying generators of the evoked activity suggests the origin of the responses to be in synaptic signaling within the nonprimary auditory area in Heschl’s gyrus or the adjacent planum temporale (Parviainen et al. 2015). The most prominent adult-age deflection at around 100 ms (N/M100) emerges surprisingly late, by the age of 9 years (Cunningham 2000; Kraus et al. 1993), although at a somewhat younger age with a longer interstimulus interval (Ceponiene et al. 2002; Takeshita et al. 2002).

Of particular interest for maturational changes and plasticity is the developmentally specific, robust response pattern in human children. In response to the passive presentation of simple sounds in children, previous studies systematically report a long-lasting and late obligatory response at around 250 ms poststimulus (referred to in the EEG/MEG literature as the M250/N250 response) (Cunningham et al. 2000; Sussman et al. 2008; Parviainen et al. 2019, which is not reported in adult studies. By this time window in adults, the obligatory activation is diminished (Johnstone et al. 1996) and activation can be measured merely related to cognitive operations such as attentional control (Huster et al. 2010). The amplitude of the developmentally specific late response has been reported to increase until about 10 years and decrease to adult values by the age of 17 (Ponton et al. 2000).Cognitively, it has been suggested to be associated with the efficiency of information processing (Ceponiene et al. 2002; Takeshita et al. 2002) both in typical (Parviainen et al. 2011) and atypical development (van Bijnen et al. 2019). However, as it is evoked in purely passive paradigms, it is likely to signify different response properties of cortical circuitry in the developing brain.

Some evidence suggests that a delayed and prolonged activation pattern can be observed in other sensory modalities of a developing brain as well. Both somatosensory cortical responses to tactile stimulation (Pihko et al. 2009) and transcranial magnetic stimulation (TMS)-evoked EEG responses in the motor cortex (Määttä et al. 2017) exhibited a sluggish late deflection in the developing brain, with similar timing to the late auditory response at 200–300 ms, in comparison to adults who evidence prominent early responses with no further distinctive activation. In general, since the later time window is more likely to reflect the stage of cortical processing related to the integration of input across sensory modalities, or top-down influence on sensory processing, it is not surprising that responses to stimuli in different modalities would exhibit similarly delayed characteristics. Hypothetically, it would make sense that developing sensory pathways demonstrate stronger and longer-lasting responsivity to the external input, benefitting the experience-driven modification of the cortical circuitry. So far, there have been no attempts to resolve whether the delayed activation pattern represents an intrinsic, species-general feature in the developing brain.

In rodent models, interest is usually in detailed cellular and laminar level functions, and it is not a common practice to measure cortical potentials, which is the usual measurement modality for the human brain. Electrophysiological responses measured from the cortex are considered a rather inexact indicator as they sum the electric activity over various events in the underlying cortical circuitry. To reach a more detailed understanding of the developmental processes in the human cortex, it is necessary to determine the correspondence in cortical phenomena between human and other mammalian (model) species that show circuitry properties similar to those of humans. Although comparisons across species have been made at the general level, the measurement protocols and techniques are typically not harmonized to enable direct parallel interpretations. Here, we used auditory stimuli to record the electrophysiological response properties of the auditory cortex in humans and rats at different ages across their developmental trajectories.

Auditory evoked responses to simple sound stimuli have been measured earlier from the mature rat’s primary auditory cortex (A1) (Barth and Di 1990; Takahashi et al. 2005). When comparing the human auditory system to other mammals, the range of frequency sensitivity or threshold levels differs, but cochlear mechanics and the generation of electrical activity function according to similar principles (Eggermont and Odenthal 1974; Kraus and McGee 1992). The smaller size of a rat’s brain results in faster conduction time and shorter latencies in response patterns compared to humans. Therefore, the well-known auditory P50 response in humans is considered to correspond to the response occurring at 10–30 ms in rats. In the same way, the counterpart of the M/N100 response is contemplated to occur at around 41–80 ms in rats, P200 at 80–130 ms, N200 at 130–200 ms, and P300 at 250–500 ms (Sambeth et al. 2003). The unquestionable benefit of evoked responses is that, when probed by simple stimuli, they give accurate information on the general timing of synaptic information transfer, with high temporal resolution. Indeed, rather than focusing on individual responses or components, it is meaningful to consider these deflections together as an evolving wave of electric activity in the cortical pathway.

It is conventional to focus on the averaged evoked responses with the underlying assumption that there is a standard, unchanged sequence of neural responses to individual stimulus presentations, and that the trial-by-trial variation can be ignored as noise. However, it may well be that—compared to the adult brain—the developing brain shows less stable timing of individual responses, which feature would not be captured by comparing average responses across ages. In fact, in the case of strong enough responses, this type of jitter would lead to the prolonged waveform in the averaged response, which has been reported as characteristic of child responses (see above, cf. N250/M250) (Parviainen et al. 2011; Ruhnau et al. 2011; Parviainen et al. 2019). An alternative explanation for the prolonged averaged response is that every single trial across the experiment shows this extended pattern of activation. The timing of synaptic signaling has been suggested as one of the core mechanisms that change along with development (Oswald and Reyes 2011), and as synaptic currents essentially generate the evoked response characteristics, it is essential to explore the single-trial characteristics of evoked responses in order to understand system-level signatures of cortical development. Specifically, clarifying the trial-by-trial stability of the prolonged response will enable us to determine whether the neurophysiological characteristics underlying the maturational changes reflect differences in the temporal precision of synaptic signaling across trials or specific, temporally distinct synaptic event(s).

This study aimed to resolve whether the late activation pattern in the auditory system is a general feature of the developing brain and can be evidenced across mammalian species. Based on the previous studies on the auditory and other sensory systems, it is reasonable to assume that the delayed juvenile-specific pattern of activity could be an evolutionarily preserved phenomenon, featuring increased excitability of the cortex by sensory stimulation. We explored whether juvenile rats and young human subjects show a similar pattern of excitability of the auditory cortex, specifically focusing on the late responses at around 250 ms poststimulus. The evoked activity is expected to be similarly delayed and prolonged in juvenile animals and children, and the amplitude is expected to decrease with age. Furthermore, we explore the trial-by-trial variance in response amplitude and latency in both species to clarify whether the prolonged response pattern reflects instability in temporal acuity or distinct and systematic synaptic events. We expect to see the age-dependent prolonged pattern at the level of single trials, ensuring that the differences relate to the prolonged reactivity of the cortical circuitry rather than imprecise timing.

## MATERIALS AND METHODS

### Subjects

#### Humans

A total of 11 adults (aged 19–26 years, 6 females) and 23 children (aged 9–13.5 years, 13 females) were recruited via university mailing lists and from schools in the Oxford area in the UK. Participants were required to speak English as their mother tongue, be right-handed, have normal hearing, and have no history of neurological diseases. Children were divided into two age groups: 9–10.5 years (group A, *n* = 11, six females) and 12–13.5 years (group B, *n* = 11, seven females). Also, 19– 26-year-old adults formed group C (*n* = 11, six females). One outlier from group B was excluded from the data since all amplitude values for all measures were 1.5 interquartile range above the third quartile. An informed consent form was collected from all adults before testing and, in the case of children, from their parents. Participants were reimbursed for the expenses incurred by their participation. The MEG measurements took place at the Oxford Centre for Human Brain Activity (OHBA) at the Warneford Hospital in Oxford. The presented study is part of a larger research project on spoken language perception in the developing brain. The study was approved by the Central University Research Ethics Committee, University of Oxford.

#### Rats

Cross-sectional rodent data from Wistar laboratory rats between 28 and 95 postnatal days was collected at the University of Jyvaskylä, Finland. Mature rats were ordered from Envigo, Denmark, and pregnant female rats were ordered from Kuopio, Finland. Juvenile rats were born and raised at the Laboratory Centre of the University of Jyvaskylä. The animals were housed in small groups in metal cages, fed ad libitum, and kept under a 12-hour light–dark cycle with lights on at 7:00 a.m. and off at 7:00 p.m. For the analysis, the rats were divided into three subgroups corresponding to the age groups of humans. Juvenile rats of age between 28 and 42 postnatal days (group A, *n* = 10, four females) were considered prepubertal (Freitas et al. 2002), juvenile rats with the age of 43–64 postnatal days (B, *n* = 10, three females) were considered adolescents (Stumpp et al. 2006), and mature rats of 65–95 postnatal days were considered young adults (Crews et al. 2000) (group C, *n* = 10, three females). See Table 1.

**Table 1.**
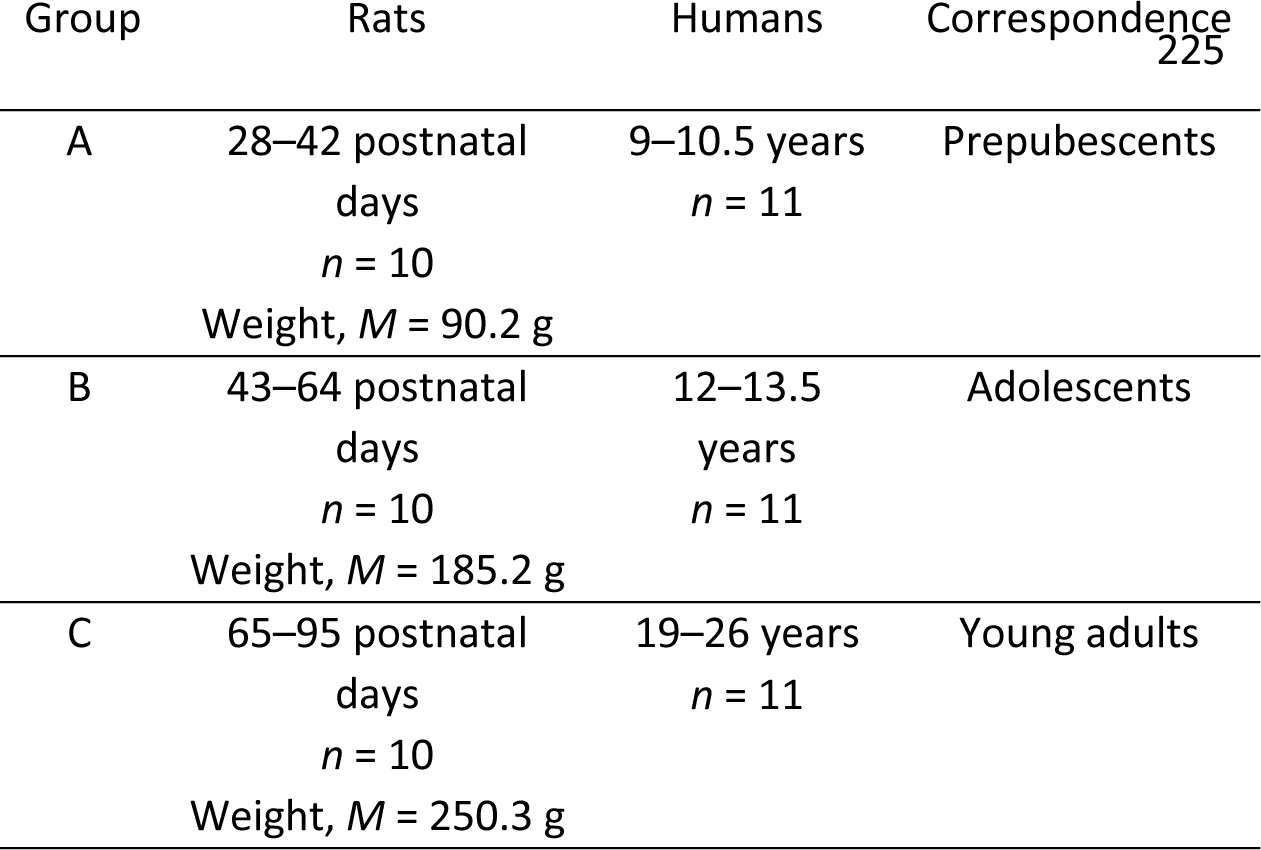
Demographic data of human subjects and rats. Age in days (rats) and in years (humans), number of individuals, mean weight (rats, in grams), and the species’ general maturational stage for the three different groups.

The experimental procedures for rodents used in this study were approved by the County Administrative Board of Southern Finland. Experiments were carried out in accordance with the European Communities Council Directive (86/609/EEC) regarding the care and use of animals for experimental procedures. All the researchers who conducted the animal experiments received special training on the care and treatment of laboratory animals, which is mandatory for using nonhuman vertebrates for research purposes. The ethical problems were discussed in terms of the three Rs (reduction, refinement, and replacement). In terms of reduction, the number of animals was adjusted so that it was possible to obtain reliable results with a minimal number of animals. In the spirit of the principle of refinement, a within-subjects design was used whenever possible. In addition, the experiments were designed so that the results answered more than one question at a time. The principle of replacement manifested in our studies as we conducted experiments in animals only when we could not answer the questions of developmental plasticity in human experiments.

### Stimuli

The aim of the cross-species comparison was to determine whether both species show a developmental difference in the excitability of the auditory cortex by simple auditory stimulation. Simple passively presented auditory stimuli were used in the recordings of both species. The stimuli were designed to be equivalent to each other in a species-specific manner. It was important to optimize the stimulation protocol for the given species, but at the same time to ensure comparability of the general response properties that are most likely influenced by temporal characteristics of stimulus presentation.

In human measurements, the stimuli were 1-kHz pure tones with a duration of 50 ms with a 15-ms rise and fall times, created in Adobe Audition 1.5 (Adobe Systems Incorporated, CA, USA). Tones were presented monaurally to the left and right ears, alternately between the ears via inserted earphones, using Presentation software (Neurobehavioral System Inc., San Francisco, CA). The interstimulus interval (ISI) was randomized between 0.8 and 1.2 s. The individual auditory threshold was obtained for each subject to ensure that all subjects had normal hearing, and during the MEG recordings, the tones were presented 65 dB above hearing level. Only right-ear sounds were used in the analysis. The stimuli in animal measurements were short clicks (duration of 1 ms) and pure tones, all presented to the right ear at 75 dB. They were 6–15 kHz sounds (duration of 100 ms with the 8-ms rise and fall times) created with the above-mentioned sound system, output from the computer sound card, and amplified with the SA1 speaker amplifier (TuckerDavis Technologies, Alachua, FL, USA). The sound intensity was measured with Bruel & Kjaer precision sound level meter type 2235 using linear weighting, peak intensity 2 kHz. Altogether, 70 click sounds were presented in one series with a random ISI of 1.8–2.2 s, and altogether 40 pure tones, frequencies of 3, 6, 9, and 15 kHz were presented in random order with the same ISI. The presentation of the sounds was programmed using E-prime, version 1.0 (Psychology Software Tools, Sharpsburg, USA). The program also generated TTL pulses to locate the start and type of each stimulus. The pulses were used to generate pulses of different amplitudes with a D/A converter to represent different stimulus types.

### MEG data acquisition and preprocessing in humans

Magnetic signals were recorded with a 306-sensor whole-head neuromagnetometer (VectorViewTM system (Elekta Neuromag oy, Helsinki, Finland) in a magnetically shielded room. Measurement sensors are arranged in triple-sensor elements at 122 locations so that every element consists of two orthogonally oriented planar gradiometers and one magnetometer. A magnetometer is a simple loop that is most sensitive to source currents that surround the loop, while a planar gradiometer with two oppositely wound in-plane loops detects the maximum signal directly above an active brain area. The position of each subject’s head relative to the recording sensors was determined using four head position indicator (HPI) coils attached to the scalp. Before the measurement, the locations of the sensors were measured with a Polhemus Fastrak® digitizer (Colchester, VT, USA). Three anatomical landmarks (nasion, left, and right preauricular reference points) were also digitized to define a head-based MEG coordinate system where the x-axis passes through the preauricular points from left to right, the y-axis passes through the nasion perpendicular to the x-axis, and the z-axis points upward. To determine the location of the HPI coils in the MEG-helmet, the coils were briefly activated at the beginning of the recordings. Using the HPI coils, it was also possible to monitor the changes in the head position during the entire measurement. Horizontal and vertical eye movements (electro-oculogram) were recorded to detect eye movements and blinks that could interfere with the data.

The MEG signals were bandpass filtered at 0.03–333 Hz and sampled at 600 Hz. The MEG data were averaged offline from 200 ms before and 800 ms after each stimulus onset. The epochs contaminated by eye movements were excluded. Only gradiometers were used, as magnetometers are more sensitive to external noise. On average, 105 (minimum 76) artifact-free trials per category were gathered from each subject. The signal space separation (SSS) method (Taulu and Simola 2006) was applied to suppress interfering signals outside of the MEG sensor array. If continuous head position indicator (cHPI) was used, SSS was applied with movement compensation. The temporal extension of SSS (tSSS) was applied when artifacts could not be removed sufficiently well with SSS. Prior to further analysis, the averaged MEG responses were baseline-corrected based on the 200-ms interval preceding the stimulus onset and low-pass filtered at 40 Hz.

### Source localization of auditory activation in humans

To localize the source currents underlying the recorded evoked responses in humans, we used ECD modeled separately for each individual (Hämäläinen et al. 1993). An ECD represents the center of activation and the mean strength and orientation of the electric current in a given brain area. As our focus was on late-emerging activation in children, the time point of maximally dipolar, distinctive field pattern within the general time window around the distinctive 250 ms-response (cf. Figure 2a) was selected for localizing the current source. In case no robust response was present in this time window, the earlier 100 ms activation peak was used for localizing activation. The 100-ms activation in adults and older children shows a highly comparable current location and orientation to the later 250-ms activation (cf. Figure 2c, d) and their topographic distribution at channel-level overlaps considerably (Figure 2e), justifying that the same macro-level model can be used to estimate the amplitude time-course of the underlying current source (see also Parviainen et al. 2011; Parviainen et al. 2019).

To determine the ECDs, a channel selection covering optimally the spatial extent of activation at the sensor level in the left hemisphere was used. As the auditory evoked activation across subjects was systematically evoked within the same general region, a standard set of 22 sensors was used for all individuals (see Figure 2e). Figure 2d illustrates the current location and orientation in the three subject groups. The source model indicated activation generally in the left temporal area, with the current orientation indicating localization to the supratemporal auditory cortex.

The strength and timing of the auditory evoked fields were further analyzed separately for each subject. In children, the most robust activation, detectable in each individual, occurred at around 250 ms after the stimulus onset (see Figures 2a and 2c). The M100 response type was distinguishable only in 3 out of 11 children in the youngest age group (A) and 2 out of 11 in the older group of children (B). In adults, the strongest response was the M100 in the early time window, and the later 250-ms activation was mostly not detectable (Figures 2a and 2c). It is worth noting that the early time window in young children is typically dominated by the P50m response, which was not quantified in this study.

### MNE source analysis

In addition to the ECD modeling, for visualization purposes and to confirm the time-course of activation independently from specific analysis algorithm, we used also minimum norm estimate (MNE)-analysis with the auditory cortex as region of interest (ROI) to the data. The data was filtered with lowpass of 40 Hz, and high pass of 0.1 Hz, blinks and heartbeat artefacts were removed using ICA and epochs of -200 to 800 ms (baseline from -200 to 0) were extracted. The auditory cortex ROI consisted of 4098 (spacing: 4.9mm, area: 24mm2) verticles at the grey-white matter boundary. For the inverse operator a loose orientation constraint (0.2) and depth weighting (of 0.8) was applied and source orientation was set to be normal to the white matter boundary, to get a more accurate representation that maintains the sign (direction of current) of the activity.

### EEG data acquisition and analysis in rodents

The EEG recordings were made in light anesthesia, known not to influence sensory processing. Corresponding to the human measures of this study, auditory responses from the left hemisphere to right ear sounds were gathered. To begin the operation, the animals were anesthetized with isoflurane (3.5%, 0.9 l/min) in a chamber. Under anesthesia, they were stereotaxically stabilized with nonpuncture ear bars, and isoflurane was applied through an inhalation mask while the concentration was reduced to 2%. Prior to the surgery, urethane was dosed intraperitoneally (1.2 g/kg, concentration 0.24 g/ml, Sigma-Aldrich, St. Louis, MO, USA), and local anesthesia (bupivacaine hydrochloridum, Bupivacaine Accord, 5 mg/ml, Orion, Finland) was applied on the surgery region to ensure the painless operation. The rat was laid on a heated platform (World Precision Instruments, Sarasota, FL, USA), and a temperature controller (ATC2000) was used with a rodent rectal temperature probe to assist in maintaining the body temperature of the animal at 38.2°C. Isoflurane was cut out 20 minutes before the acute recordings. During the measurements, the depth of anesthesia was controlled by regular testing of the withdrawal pedal reflex. Supplementing doses of urethane (approx. 0.1–0.2 ml) were applied during the operation if needed to maintain a proper level of anesthesia. Also, 2 ml of saline was added every two hours for adult rats and around 0.5 ml for juvenile rats to maintain a stable physiological condition.

For the auditory epidural recordings, a craniotomy was conducted. The skin and muscles over the horizontally located skull and the left auditory cortex were removed. The skull was cleaned with cotton buds and hydrogen peroxide and washed with saline. If needed, the bleeding of the muscles was stopped using silver nitrate. The exact location of the primary auditory cortex (A1) was calculated by using the anatomical point bregma as a landmark. In adults, an opening was drilled to the squamosal bone of the skull: from the bregma anterior-posterior, (−4.5)–(−6.5) mm; and dorsoventral, 3–5 mm. In juveniles, the size of the opening was adjusted according to the age of the rat concerning the distance between the bregma and lambda. Two holes were drilled over the frontal cortex, and screws (Ø 1.0 mm) were driven to the bone. Screws were attached to the end of a 10*80 mm Plexiglas plate with acrylic, and the other end of the plate was fixed to the stereotaxic instrument.

For the reference electrode (a 28-gauge stainless steel needle, Ø 0.36 mm, BD Lo-Dose syringe, USA), a 1-mm diameter hole was drilled over the right cerebellum 1.5 mm posterior to the lambda and 2 mm lateral to the midline. The needle was inserted into the hole 1 mm below the surface of the cerebellum and fixed to the skull with cyanoacrylate glue. The ground electrode was a needle similar to the reference electrode and inserted subcutaneously in the neck. A silver wire recording electrode (Ø 0.5 mm, A-M Systems, WA, USA) was positioned on the surface of the dura and attached to a miniature connector together with ground and reference electrodes. The right ear bar was removed, and a speaker (MF1 magnetic speaker, Tucker-Davis Technologies, Alachua, FL, USA) was positioned at the level of the ear and toward it at a distance of 20 cm. After the bar was removed, the head remained stable with the previously fixed acrylic plate.

The recordings were performed in a Faraday cage. A short stimulus test series of click sounds with an ISI of 2 s was played before the acute recordings to ensure that the techniques work properly and that the area of measurement was valid. Large veins were avoided when the electrode was placed. The electrode was positioned over the different parts of the opening, and the location of the largest positive response at around 50 ms poststimulus was selected as the position of recording. This positive deflection indicates the synaptic responses to thalamic input to the auditory cortex, and thus the tone with the best representation at the recording site. At each site, 3–5 responses were recorded to evaluate the amplitude of the response. During the recordings, spontaneous and evoked neural activity was monitored online. The signals were initially amplified 10x with the MPA8I head stage and then high-pass filtered at 0.1 Hz, amplified 50x, and low-pass filtered at 5 kHz with an FA16I filter amplifier (all from Multichannelsystems, Reutlingen, Germany). The signals were digitized using an ME64 system (Multichannelsystems) at 20 kHz, low-pass filtered to 600 Hz, downsampled to 2 kHz, and written to a hard disk using the MC_Rack program (V 4.6.2, Multichannelsystems). After the recordings, the anesthetized animals were administered an overdose of general anesthetics by i.p. injection (urethane, Sigma-Aldrich, St. Louis, MO, USA).

The EEG data were analyzed with the BrainVision Analyzer 2.1 program (BrainProducts, Gilching, Germany). Epochs 200 ms before and 800 ms after the stimulus were segmented, visually monitored for artifacts and averaged for each stimulus type. However, no sweeps were rejected from any animals, and no animals were excluded from the final analysis. The averages were written into text files, imported to Excel, and averaged over animals in each age group.

The stimulation paradigm in rats was harmonized with the human experiment in terms of ISI, but the other stimulation parameters (frequency, sound type) were adjusted to achieve maximally reliable responses in rats. Because the click sounds evoked better-quality responses, they were used for statistical analysis instead of sine-wave tones. However, sine-wave tones (which were used in humans) gave essentially identical general-level findings. For comparison, Appendix A reports the average response to sine-wave tones in rats.

### Single-trial analysis in humans and rodents

Besides the analysis of averaged responses in each individual, the single trials, i.e., the epochs of electric activation evoked by each sound stimulus, were also analyzed in each subject. Figure 1 shows a pipeline of single-trial analysis in humans and rats. In humans, for extracting single trials, the continuous time-series of the original data was first reproduced at the source level by applying the ECDs obtained from the averaged data by using MNE-Python functions. The single-trial time series were epoched from the continuous data for further quantification of the trial-to-trial consistency of the evoked response amplitude in different time windows. For the electrophysiological recordings of the rat auditory cortex, continuous recording was used directly to extract single-trial epochs.

**Figure 1.**
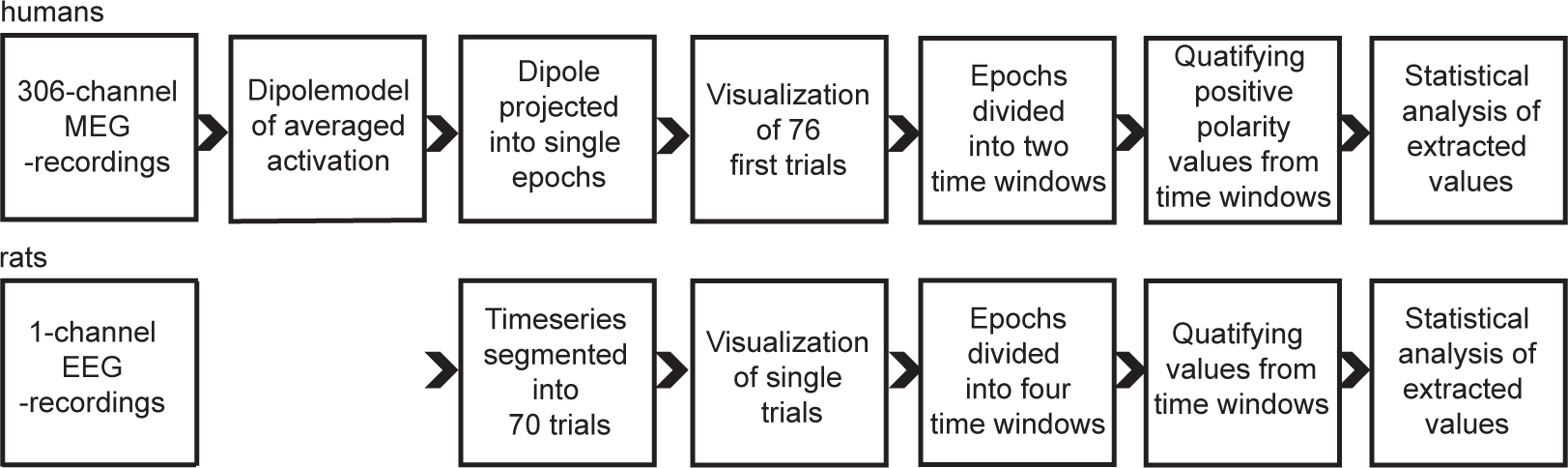
The single-trial analysis pipeline. Analysis steps for extracting the single-trial latency and amplitude values for the MEG recordings in humans and EEG recordings in rats.

For analyzing the trial-to-trial stability of the local maxima in the amplitude waveform, the time course of auditory epochs was divided into early (0–150 ms) and late (150–450 ms) time windows in humans and into three early (0–10 ms, 10–30 ms, and 30–150 ms) and one late (150–450 ms) time windows in rats. The first 0–10-ms time window in rats covers the fast brainstem responses, and the following 10–30-ms time window reflects the peaks originating in subcortical and early sensory cortical circuits. In humans, the same stimulation settings cannot be used to produce reliable evoked responses both from the brainstem and subcortical or primary cortical structures at the early time window (0–50 ms) and from nonprimary auditory areas at the later (>50 ms) time window. In other words, brainstem responses, middle-latency responses, and late cortical components all require specific ISI and stimulus properties. In this study, we were mainly interested in the development of the late cortical activation, and thus the stimulus paradigm was optimized for the late cortical responses. Therefore, the amplitude values are not extracted for the earliest components in humans (i.e., middle latency responses and brainstem responses). All 70 measured epochs were analyzed in animal subjects. In humans, the final number of artifact-free epochs varied, and based on the minimum number of trials in individual subjects, the first 76 epochs were analyzed in all human subjects.

### Statistics

#### Averaged waveforms

To analyze the averaged waveforms in humans, we collected both amplitude and latency measures (maximum amplitude and peak latency) separately for each individual in the early (50–150 ms, M100) and late (150–350, M250) time windows (cf. Figure 2). In rats, the same activation measures were collected separately from the following time windows: 0–12 ms (N6), 0–25 ms (P14), 12–37 ms (N21), 25–75 ms (P41), 50–150 ms (N108), and 150–350 ms (P214). See Table 2.

**Table 2.** Time windows (ms) of analyses. Time windows were used for extracting maximum amplitudes and peak latencies for the statistical analysis of single-trial and averaged data in humans and rats.

| Response level | Rats |  | Humans |
| --- | --- | --- | --- |
|  | Single trials | Averages | Single trials and averages |
| Brainstem | 0–10 | 0–12 | - |
| Early cortical I | 10–30 | 0–25<br>12–37 | - |
| Early cortical II | 30–150 | 25–75<br>50–150 | 0–150 |
| Late cortical | 150–450 | 150–350 | 150–450 |

A one-way analysis of variance (ANOVA) was used to evaluate the age differences in the strength of averaged auditory responses separately in humans and animals. A post hoc test with Bonferroni correction was used to compare the means of the three age groups. The alpha level was *p* < .05. Levene’s test of equality of error variance indicated that all of the groups had the same variance.

#### Single trials

For single-trial analysis, the maximum values of evoked deflections were collected from time windows 0–150 ms and 150–450 ms in humans and from time windows 0–10 ms, 10–30 ms, 30–150 ms, and 150–450 ms in rats. In addition, the peak latencies were collected by computing the time points at which the maximum values were reached in each time window. If the activation level was below zero during the time window of a positive deflection, the epoch was cut out of the analysis. Similarly, if the activation remained over zero during the time window of negative deflections, the epoch was excluded. The definitive number of single epochs differed between individuals and age groups, mostly when individual variation was high. Appendix B shows the number (*N*) of single epochs in each response type and age group. Human data were based on dipole-modeled time series, and the source model (ECD) represents the current with a certain location/direction within the cortical laminae. Therefore, we did not use the amplitude waveform to extract values from the “negative” part, as we cannot be sure that the source estimate correctly captures the activation within these time points— and the results are not reliable.

Multilevel multigroup analysis with pairwise comparisons (Mplus version 8.4) was used to analyze the statistical data of single trials. The variance in peak latency and amplitude differences between age groups and time windows were tested with the Wald test, and further comparisons were conducted pairwise. The maximum likelihood estimation method with full information was used (FIML). All the available information was included in the analyses, and missing data were assumed to be missing at random (MAR). Within-level variation reflected *internal variation* of the subject’s (human or rat) single-trial responses, and between-level variation reflected *between-subjects variation* of responses. Since intraclass correlation (hierarchy within the data, i.e., the degree to which between-subjects variation accounts for total variance) was detected, a two-level test was conducted for the latency values. Mean differences of maximum amplitudes in the different time windows across the three age groups were tested with a complex method modeled at a single level (time window), yet the hierarchy of the data organization was taken into account by correcting standard error and *p*-value. The hierarchy (i.e., the fact that each subject has multiple values and the data contains multiple subjects) of the data was based on a cluster (i.e., subject).

## RESULTS

Figure 2a depicts the time course of the sound-evoked activation (auditory evoked response) in one sensor in the left hemisphere of each age group. For comparison, Figure 2b shows the auditory evoked response in the three age groups of rats, as measured with the silver wire electrode from the surface of the primary auditory cortex. The time course indicates two separate time windows of activity, at around 100 ms and around 250 ms. Figure 2c visualizes the time course of activation in the auditory cortex ROI in the left hemisphere, evidencing the robust late-emerging 250-ms activation in the child groups. The ECD model of the source current in each age group, and the distribution of sensors that were used for their localization are depicted in Figure 2 d and 2 e.

**Figure 2.**
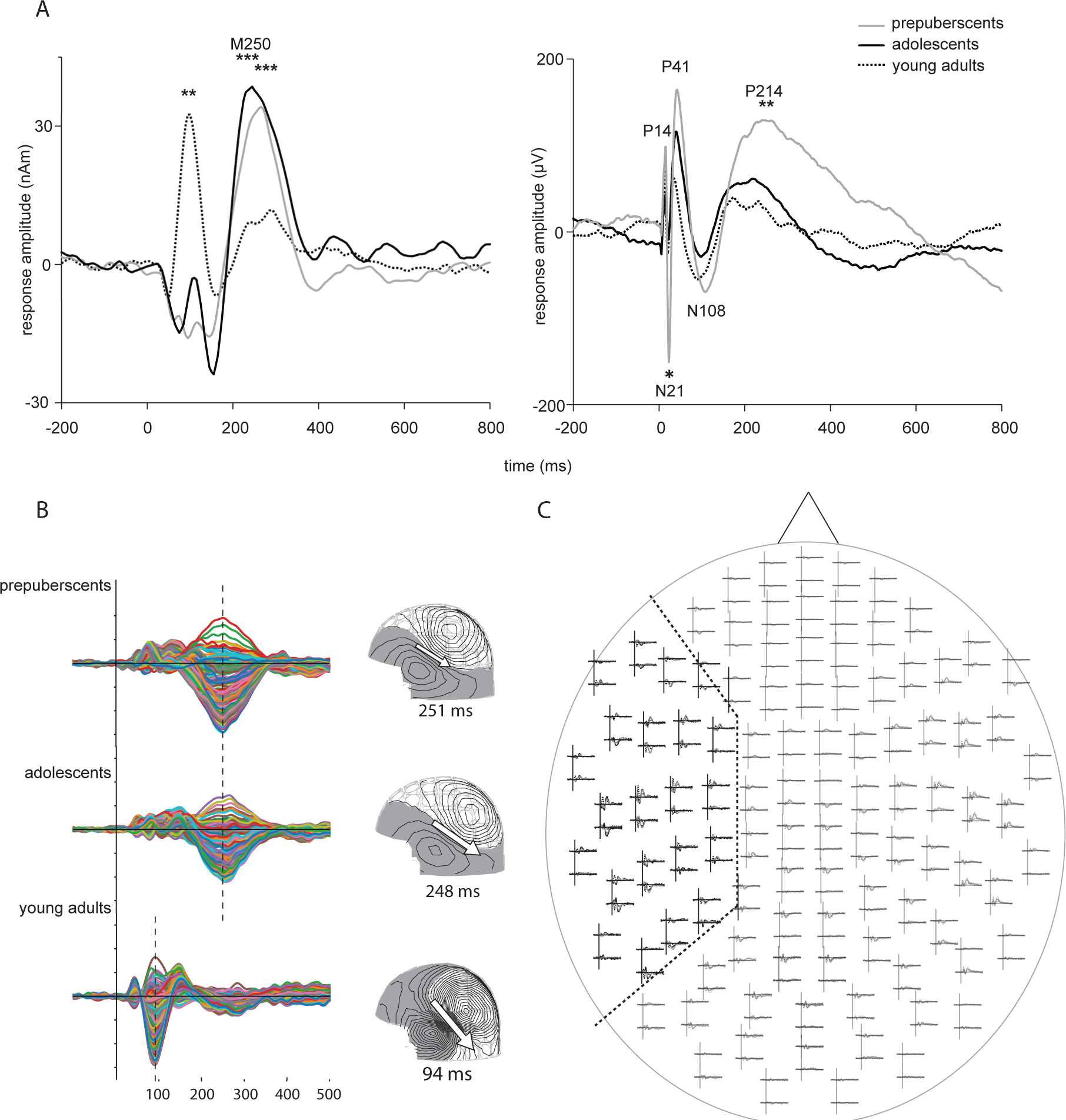
Time course and location of auditory activation in humans and rats. (a) Time course of activation evoked by the auditory probes in the current source located in the left supratemporal auditory cortex in humans. (b) Time course of activation evoked by the auditory probes as recorded by the silver wire electrode from the surface of the auditory cortex in rats. (c) Butterfly plots illustrating the time course of evoked activity in individual vertices within the left supratemporal auditory cortex in humans. (d) The location and orientation of the equivalent current dipole (ECD) in humans, fitted at the time point of maximum amplitude for each age group (250 ms for children, 100 ms for adults). (e) Whole-head sensor array with evoked responses in the three age groups of humans. The distribution of sensors used for localizing the ECD is delimited with a dashed line.

### Late auditory evoked activation is similarly delayed in children and juvenile rodents

#### Late auditory activation pattern in humans across different age groups

The late auditory response (M250), extracted from the time window of 150–350 poststimulus, peaked on average at 284 ms (latency range for maximum: 193–325 ms across all the human subjects). The main effect of age was evident in the amplitude of the response (maximum, (F(2, 30) = 29.85, *p* < .001)), and children showed a significantly stronger response in this time window than adults (A vs. C, *p* < .001; B vs. C, *p* < .001).

#### Late auditory activation pattern in rats across different age groups

The late auditory response (P214) occurred on average at 214 ms in rats (latency range for maximum: 150–338 ms across all animals). As in humans, the main effect of age was evident in the amplitude of the response (maximum, (F(2, 27) = 7.95, *p* < .01), and there were significantly stronger responses in prepubertal rats than in mature rats (A vs. C, *p* < .01) and in prepubertal rats than in adolescent rats (A vs. B, *p* < .01). Furthermore, the main effect of age in the peak latency of the P214 response (F(2, 27) = 6.95, *p* < .01) reflected longer response latencies in the prepubertal rats (247 ms) than in the adolescent (194 ms) and adult rats (202 ms) (A vs. B, *p* < .01; A vs. C, *p* < .05).

### Early transient response pattern shows species-specific differences

#### Early auditory activation in humans

The early auditory response component peaked on average at 73 ms (M100) in humans (at a latency range of 30–98 ms) (see Figure 2a) and showed the main effect of age group in amplitude (maximum, (F(2, 30) = 16.02, *p* < .001): Adults had significantly stronger M100 responses than prepubertal children (C vs. A, *p* < .001) or adolescents (C vs. B, *p* < .001).

#### Early auditory activation pattern in rats

The early auditory response pattern before the prolonged late response comprised three positive and three negative deflections in the rats (see Figure 2b). The first distinguishable component at around 6 ms (N6, at a latency range of 3–7 ms across all animals) or the second component at around 14 ms (P14, at a latency range of 10–21 ms across all animals) showed no differences between the age groups in the maximum amplitude. The following component at around 21 ms (N21, at a latency range of 13– 26 ms across all animals) showed a main effect of group in response amplitude (F(2, 27) = 8.50, *p* =.001), being stronger in the prepubertal rats than in both the adolescent and adult rats (A vs. B, *p* < .01; A vs. C, *p* < .01). In the following components at around 41 ms (P41, at a latency range of 34–50 ms across all animals) and at around 108 ms (N108, at a latency range of 75–144 ms), there was no effect of age on the amplitude. In sum, only the 21-ms response evidenced age-group differences, with stronger amplitude in the youngest age group than in the two older ones.

### Juvenile pattern of auditory evoked responses is distinguishable at the level of single trials in both humans and rats

Figure 3 depicts the single-trial auditory evoked responses for three human and rat individuals, one from each age group (see Appendix C for all individuals). Visualization of the individual epochs shows a similar emphasis on activation in the late time window (at around 250 ms) in young individuals (uppermost panels), as was evident in the average responses. In adult/mature individuals, on the other hand, activation at the early stage (at around 100 ms) seems to show a more systematic response than later activation. For statistical analysis of age-related differences, the maximum values of evoked deflections (maximum amplitude) and the time points when the maximum values were reached (peak latencies) were collected from time windows 0–150 ms and 150–450 ms in humans and from time windows 0–10 ms, 10–30 ms, 30–150 ms, and 150–450 ms in rats. In humans, the single-trial amplitude reflects the amplitude of current with the specific orientation (cf. Figure 2b), while in rats, the amplitudes reflect the electric potential difference between the measurement electrode and the reference.

**Figure 3.**
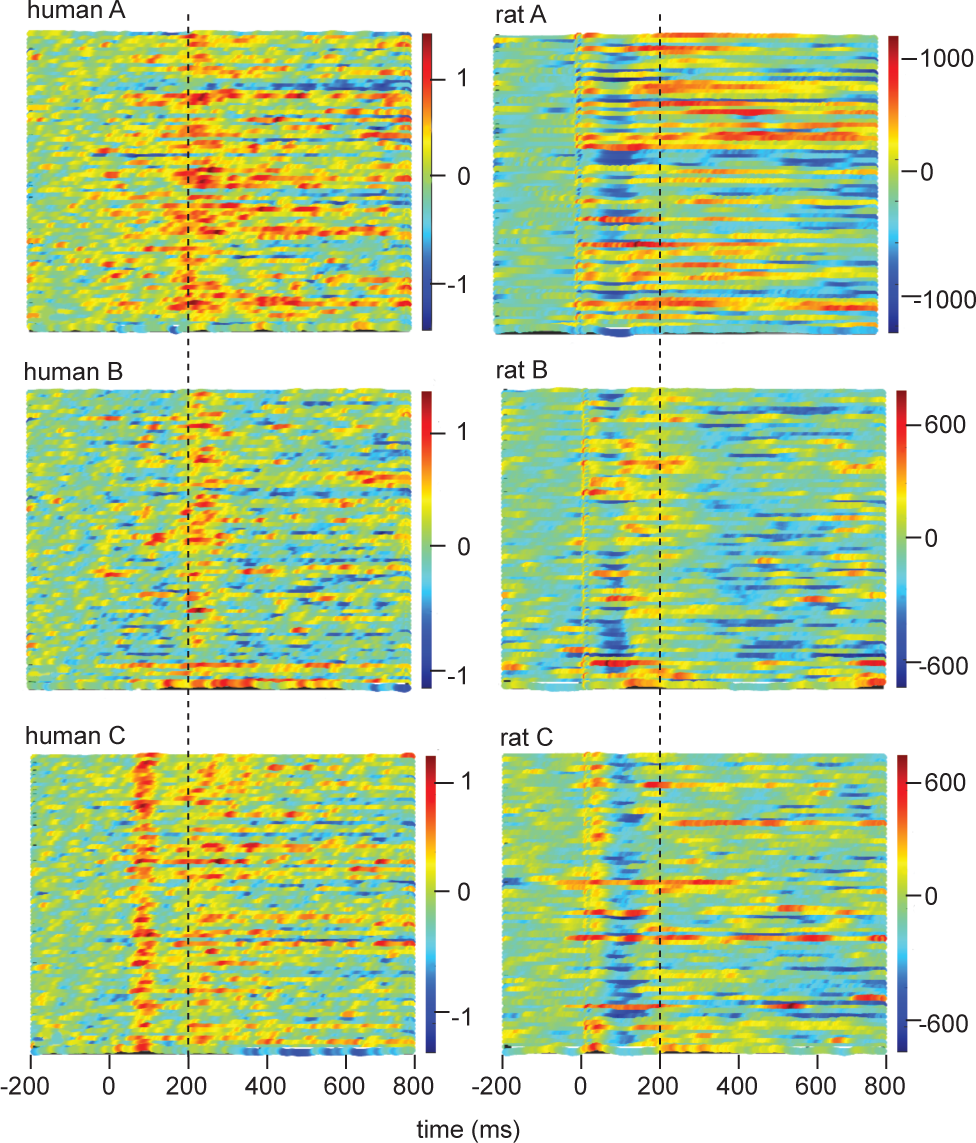
Time-varying amplitudes in single trials in humans and rats. Single-trial amplitude time series of individual subjects’ auditory responses. The response amplitude values are illustrated with color codes across time in the horizontal direction, and consecutive trials are run in vertical columns starting from the first trial in the uppermost part of each subfigure. Dashed lines depict the time windows used for collecting amplitude and latency values for the single-trial analysis.

The variance in peak latencies across different age groups was studied by multilevel multigroup analysis with pairwise comparisons (Mplus version 8.4). Both within-level (within-group) variation (i.e., *internal variation* of the human or rat single-trial responses) and between-level (between-group) variation (i.e., *between-subjects variation* of responses) were tested. Differences in amplitudes across the different time windows, separately in the three age groups, were tested with a complex method, modeled as a single level (time window) but accounting for the hierarchy (i.e., each subject has multiple values and data contains multiple subjects) of the data based on a cluster (subject).

#### Single-trial timing in humans: responses in the late time window show higher stability in children than in adults

Figure 4 demonstrates the distribution of the single-trial peak latencies in the two time windows in humans. The intraclass correlations (reflecting the reliability of the clustered structure in the data) were significant both at the late (9%, *p* = .05) and at the early (11%, *p* < .01) time windows, indicating that part of the total variance was accounted for by between-subjects variation, i.e., groups.

**Figure 4.**
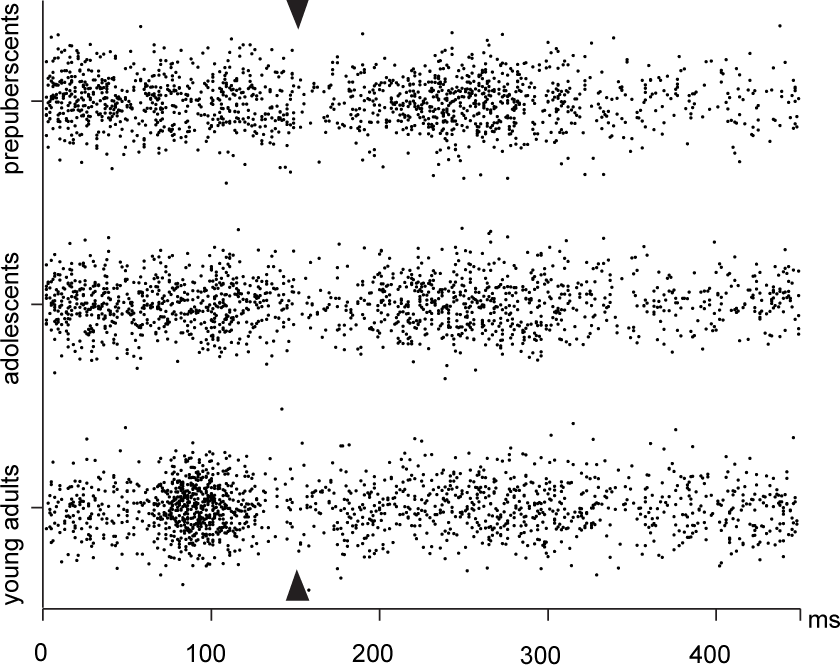
Timing of activation peaks in single trials in humans. The latency distribution of the local amplitude maxima of the single trial evoked responses in humans, extracted in two time windows, early (0–150 ms) and late (150–450 ms). The arrows point to the cut point between the time windows (150 ms).

In the late time window, the total variance (i.e., including variance at within and between levels) of peak latency values differed significantly by age group (W(2) = 17.93, *p* < .001). The variance in the prepubescents (A) was smaller than that in the adolescents (B) (*p* < .05) and in adults (C) (*p* < .001). When this total variance was examined at within (i.e., single-trial variance in each individual) and between (i.e., variance between individuals) levels, the age groups differed for within-level comparison (W(2) = 5.93, *p* < .05): individuals in group A had a smaller variance across single-trial values than individuals in group B (*p* < .05). However, the groups did not differ in between-level comparison.

Also, in the early time window, the total variance of peak latencies differed significantly by age group (W(2) = 14.04, *p* < .001). The total variance was smaller in adults (C) compared to prepubescents (A) (*p* < .001) and adolescents (B) (*p* < .001). In between and within levels, comparisons were insignificant. In sum, the most stable and systematic activation emerged at around 100 ms in adults and around 250 ms in young children, which is evident in Figure 4 as more densely distributed values for prepubescents than for adults in the late time window and the opposite in the early time window.

#### Single-trial timing in rats: age-related differences in single-trial stability are less systematic in rats than in humans

Figure 5a shows the peak latency distribution in the rats. Similar to humans, the local maxima identified in the average waveform (N6, P14, N21, P41, N108, and P214) were also evident at the level of single-trial distribution in the four time windows. Figure 5b illustrates the early peak latencies of rats on a more accurate scale. Overall, the emphasis on the auditory response pattern seemed to shift to earlier latencies along with maturation (see the peaks at 41 ms, 21 ms, 14 ms, and 6 ms).

**Figure 5.**
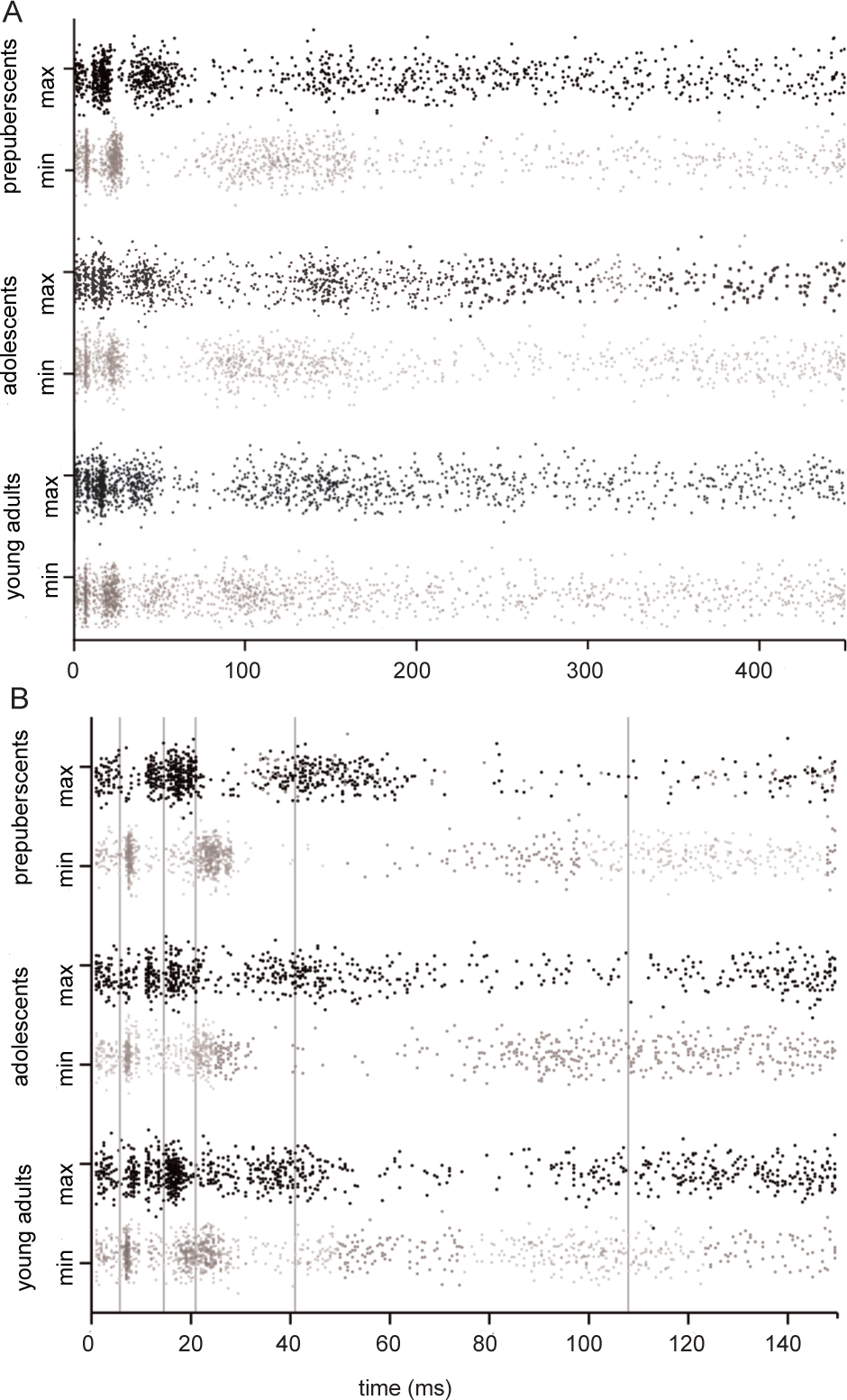
Timing of activation peaks in single trials in rats. (a) The latency distribution of the local amplitude maxima of the single-trial evoked responses in rats, extracted from time windows 0–12 ms, 0–25 ms, 12–37 ms, 25–75 ms, 50–150 ms, and 150–350 ms. (b) The latency distribution of early responses in rats by age groups expanded from Figure 5a.

Out of the deflections with negative polarity, intraclass correlations were significant for the peaks at 10–30 ms (44%, *p* < .001) and at 30–150 ms (14%, *p* < .01), but not for the peak at 0–10 ms (9%, *p* = .11). Out of the deflections with positive polarity, intraclass correlations were significant for the peaks at 10–30 ms (33%, *p* < .001) and at 30–150 ms (27%, *p* = .001), but not for the peak at 150–450 ms (4%, *p* = .10).

The total variance of peak latency values at the late response (P214) did not differ significantly by age group (W = 5.5, *p* = .06); differences in within and between levels were also not significant. With a *p*-value close to the significance level, the absolute level of latency variance was smaller in the young groups than in the adult group (similar to the human data).

The variance of peak latency values at earlier responses was inspected individually in the rats. The P14 response did not differ in total or within levels, but in between-level variance, there was a significant difference (W = 9.44, *p* < .001). In pairwise comparison, group C showed a larger variance than group B (*p* < .01). The P41 response did not have a variance difference in total, yet in within-level variance, there was a significant difference (W(2) = 6.35, *p* < .05): the variance across single trials within individuals was larger in group B than in group A (*p* < .05). In addition, the between-level variance showed a significant difference (W(2) = 10.12, *p* < .01), and group C showed a larger variance than group B (*p* < .01).

For the N6 response, there were no total, within-, or between-level differences in the variance of the peak latency values. For the N21 response, the variance at the total or between level did not differ. At the within level, there was a significant group difference (W(2) = 10.72, *p* < .01): in group B, the internal variance was larger than in group A (*p* = .001). The N108 response latency showed a significant variance difference in total (W(2) = 6.02, *p* < .05), and pairwise comparisons indicated that there was significantly larger variance in group C than in group A (*p* < .05). Also in between-level variance, the difference was significant (W(2) = 7.2, *p* < .05): group C had a larger variance than group A (*p* < .01) or B (*p* = .01). There were no significant within-level differences.

In sum, the peak latency values in rats showed more instability than in humans. No consistent developmental differences in rats emerged; however, some pairwise comparisons indicated larger variances in adult group C.

#### Single-trial amplitude variance in humans: unlike adults, children show stronger emphasis on long-latency than on early-latency responses

Figure 6 illustrates the results of the complex analysis of the amplitude differences between the time windows in the three age groups of humans. Although a similar sequence of auditory responses was consistently demonstrated in subjects across all age groups (encompassing early and late peaks), the emphasis of activation across the time windows seemed to depend on age. The complex method showed significant amplitude differences between time windows (W(2) = 36.58, *p* < .001). The single-trial evoked responses were stronger in the late time window (150–450 ms) than in the early time window (0–150 ms) in child group A (*p* < .001) and child group B (*p* < .001). However, in adult group C, there was no difference between the early and late time windows in single-trial peak amplitudes.

**Figure 6.**
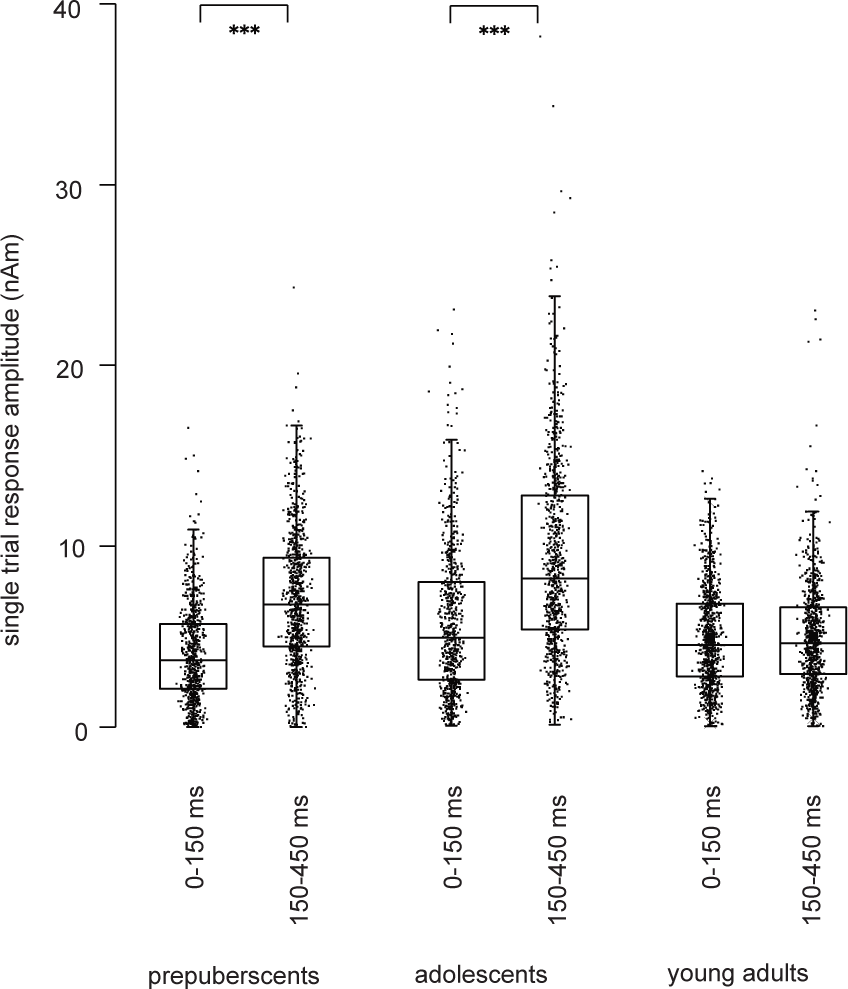
Amplitude of activation peaks in single trials in humans. Maximum amplitude values of individual responses in humans in the early (0–150 ms) and late (150–350) time windows.

#### Single-trial amplitude variance in rats: unlike mature rats, juvenile rats show stronger emphasis on long-latency P214 response than on preceding peaks

Figure 7 illustrates the results of the complex analysis of the amplitude differences between the time windows in the three age groups of rats, modeled at a single level. The analysis of the single-trial amplitude values in rats revealed essentially similar results to the comparative analysis in humans.

**Figure 7.**
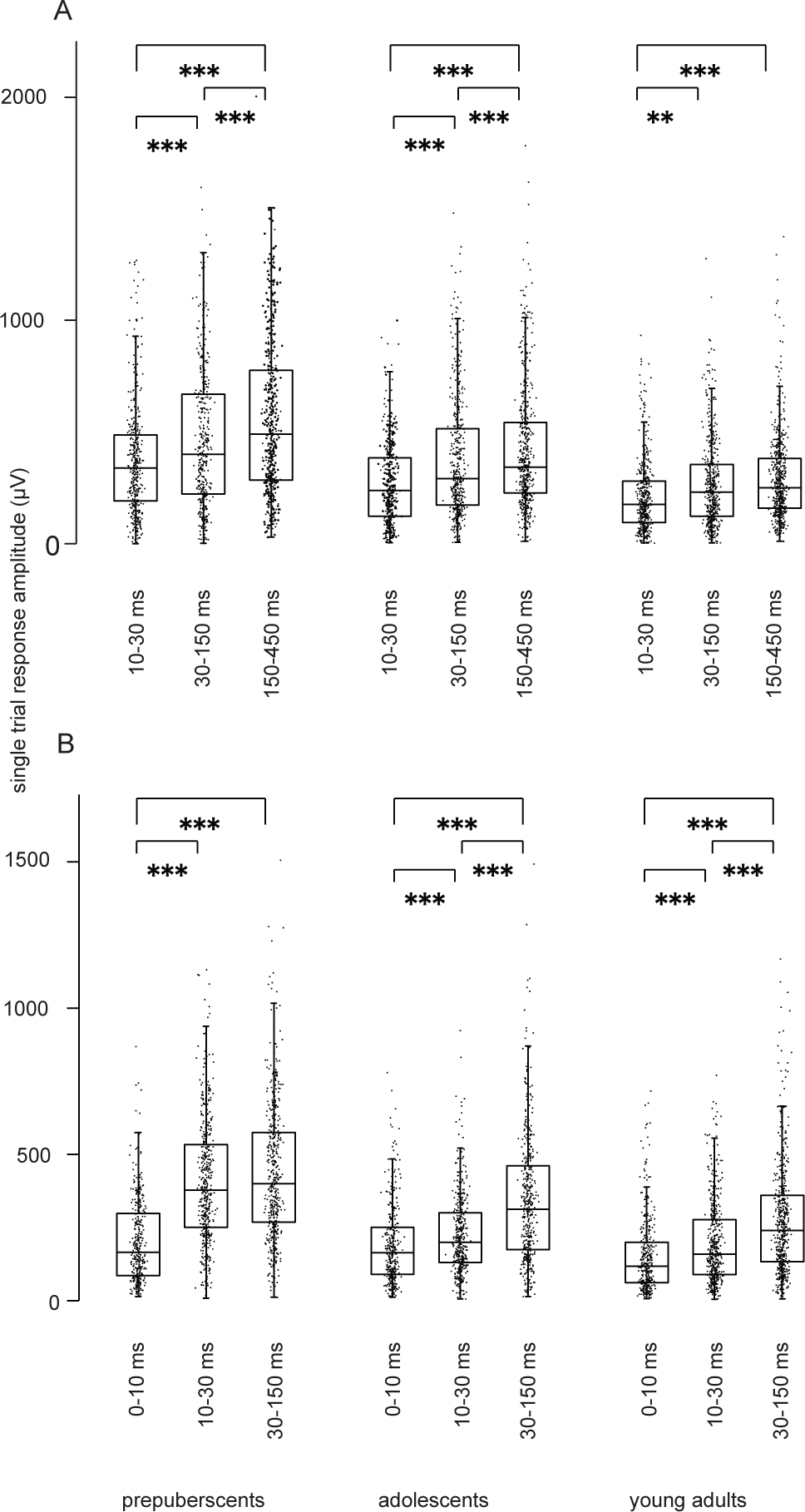
Amplitude of activation peaks in single trials in rats. (a) Single-trial maximum values for positive peaks, i.e., for P14, P41, and P214, separately in each age group of rats. (b) Single-trial maximum values for negative peaks, i.e., for N6, N21, and N108, separately in each age group of rats.

The maximum single-trial amplitudes for P14, P41, and P214 (Figure 7a) showed in general significant differences (W(2) = 21.81, *p* < .001). In youngest group A, the single-trial peak amplitudes across the three time windows differed significantly (W(2) = 39.02, *p* < .001): amplitude values were larger in the third (P214) than in the first (P14) time window (*p* < .001) or in the second (P41) time window (*p* < .001) and also larger in the second than in the first time window (*p* = .001). Adolescent rats (group B) also showed significant differences (W(2) = 22.73, *p* < .001). The amplitudes were larger in the third than in the first time window (*p* < .001), in the third than in the second time window (*p* < .001), and in the second than in the first time window (*p* < .001). Mature rats in group C had significant differences as well (W(2) = 54.89, *p* < .001): the amplitudes were larger in the second than in the first time window (*p* = .01), in the third than in the first time window (*p* < .001), but not in the third than in the second time window.

Analyzing single-trial maximum amplitudes of N6, N21, and N108 showed significant differences (W(2) = 40.45, *p* < .001) (Figure 7b). In group A, a single-level test for the time window showed significant differences (W(2) = 165.35, *p* < .001), and the amplitudes of the responses were larger in the third than in the first time window (*p* < .001) and in the second than in the first time window (*p* < .001). There were no differences between the amplitudes in the second and third time windows. In group B, significant differences were shown (W(2) = 75.6, *p* < .001), and amplitudes in the third and second time windows were larger than in the first time window (3 vs. 1, *p* < .001; 2 vs. 1, *p* < .001), and amplitudes in the third time window were larger than in the second time window (*p* < .001). In group C, significant differences were also shown (W(2) = 59.22, *p* < .001), and comparisons by time windows showed similar results to those in adolescent group B. The amplitudes were larger in the second and third time windows than in the first (2 vs. 1, *p* < .001; 3 vs. 1, *p* < .001) and in the third time window than in the second (*p* = .001).

In summary, the time window differences of single-trial amplitudes confirmed the finding of averaged waveforms, showing that the P214-ms response amplitudes compared to the preceding response amplitudes were systematically pronounced in young rats. In group A, the amplitudes of the N21-ms and N108-ms did not differ, while the older groups had smaller N21-ms amplitudes compared to N108-ms amplitudes.

## DISCUSSION

We aimed to clarify whether a delayed and prolonged pattern of evoked activity is characteristic of the juvenile auditory cortex across species. Our findings demonstrate that human children and young rodents exhibit similarly emphasized long-latency response at around 200–250 ms probed by simple auditory stimuli, whereas in the mature auditory cortex the late activation is diminished both in humans and in rodents. We further showed that the juvenile-specific delayed responsivity of the cortex is evident at the level of single trials of young individuals. We interpret this robust prolonged electric current to demonstrate the increased excitability of the cortical synaptic circuits.

Given the long phylogenetic distance, the age-related differences in the late cortical responses to auditory stimuli were surprisingly similar in humans and rats. Prepubertal rats exhibited stronger and temporally more extended late responses than mature rats, with remarkably similar absolute timing compared to the human responses. Unlike in humans, in rats, the response was stronger in prepubertal rats than in adolescent rats. This may be due to the faster maturational trajectory in rodents than in humans, even though we aimed at matching the maturational stage. Given the differences in the higher cognitive functions and, for example, in means of communication, this shared characteristic of the developing cortex presumably reflects a fundamental property of neural signaling that is meaningful for the developing nervous system.

In humans, a prolonged activation pattern has also been reported earlier (Ponton et al. 2000; Sussman et al. 2008; Parviainen et al. 2019;), but in electrophysiological recordings of rodents, the focus has been on the early responses, and the epoch of analysis is typically limited to the first tens of milliseconds. This is justified in studies of passive sensory processing, as the neural signal progresses rapidly beyond primary auditory areas. Indeed, after 100 ms, the evoked activation is usually associated with cognitive aspects, sensory integration, or top-down processing of the stimuli that are less frequently approached in nonhuman species. This convention may, however, leave unnoticed developmentally meaningful slower sensory responses. Although the sensory pathway transmits the signal rapidly to higher-level areas in the juvenile brain, it may well be that, in parallel, developing sensory cortices exhibit recurrent synaptic activity. This type of prolonged electric discharge may be important to enable the formation of associations (hence learning) within a regional and especially cortico-cortical network, therefore indicating the level of cortical plasticity.

The emphasis on late versus early activation appeared rather systematic across individual trials in the juvenile cortex. In human children and adolescents, the single-trial amplitudes were consistently stronger in the late (>200 ms) than in the early (<200 ms) time window, and both prepubertal and adolescent rats showed systematically emphasized amplitudes for the responses in long-latency versus middle-latency time windows (i.e., P214 vs. P41). This difference between time windows in the single-trial response amplitudes seemed to vanish along with development and was not present in mature subjects (either in humans or in rats).

In addition to the emphasized amplitude of the late M250/P214 response in juvenile individuals, the single-trial analysis in humans indicated age-dependent differences in the trial-by-trial variation in the timing of activation. The variation in M250 response maximum latencies, in other words, the “jitter” of the responses, appeared smaller in prepubertal children than in adolescents or adults. In rats, the maturational differences in trial-by-trial variance in response latency for the late response were not that explicit. Although the absolute level of jitter in P214 latency was similarly smaller in juvenile rats than in human children, no statistically significant group differences emerged.

The smaller (or equal) temporal variance in younger than in mature individuals contradicts one of our two hypothesized explanations underlying the prolonged response pattern in the juvenile brain, namely the increased trial-by-trial variation in timing. It seems unlikely that volatile temporal precision in synaptic information processing would explain this response. On the contrary, when probed by simple auditory stimuli, the juvenile auditory cortex seems to systematically exhibit a prominent, sluggish response, likely reflecting distinct synaptic signaling and the unique response property of the juvenile cortex.

Passively presented sounds can be considered “probes” of cortical excitability (by external sensory information). Thus, the prolonged response in children could indicate increased excitability of the cortical circuitry at these long latencies. These juvenile stage responses may indicate decreased levels of inhibition, perhaps due to the well-established protracted developmental trajectory of inhibitory GABAergic synaptic signaling (Le Magueresse and Monyer 2013), specifically demonstrated for the auditory cortex by Takesian et al. (2010). Interestingly, while in the averaged analysis the difference between age groups in early vs. late activation seemed robust (adults show stronger activation in the early than in the late time window, whereas for children the pattern is opposite), in the single trials this difference is somewhat less clear. Indeed, it seems that, especially in rats, mature individuals occasionally show responses in the late time window.

This finding may be related to the fluctuation in the ongoing brain states that is likely to influence evoked activity (Civillico and Contreras 2012). Unlike humans, rats were measured in light urethane anesthesia, roughly corresponding to the human sleep stage, which has been associated with increased N2 amplitudes in response to sounds (Nielsen-Bohlman 1991). Sleep, especially the non-REM sleep stage, is characteristically associated with endogenous slow waves (SW) that reflect transient increases and decreases in cortical excitability. Timofeev et al. (2020). speculated that these SWs could relate to the process of modifying network connectivity in support of learning, also in the developing brain. If our externally evoked slow response in the 200–300-ms time window links with similar processes, it could be interpreted to signify the level of inhibitory synaptic signaling (or lack thereof) in the cortical sensory circuitry. Indeed, in the developing brain, it is beneficial to allow a burst of recurrent activation in the sensory cortices that increases the likelihood of simultaneous synaptic signaling (i.e., Hebbian learning).

The early transient response components showed stronger species-specific differences, despite the similar experimental paradigm across species. This was to be expected from the extensive literature on auditory cortical responses in each species. In humans, in line with previous studies (Hillyard and Picton 1987); Parviainen et al. 2005), adults exhibited a prominent M100 response, which was less clear in younger groups. Interestingly, the single-trial analysis in humans revealed that the temporal consistency of activation (i.e., the intertrial precision of signaling) in the M100 response was also higher in adults than in children.

In rats, the higher signal-to-noise ratio (and focal measurement from the brain surface) enabled the identification of the more detailed canonical sequence of auditory evoked responses before the prolonged pattern. These early responses, starting from the brainstem auditory evoked potential (BAEP) at 6 ms, were remarkably robust even at the single-trial level. Overall, the early responses seemed to reach mature characteristics at an earlier age than the late prolonged pattern since there were no differences in the amplitudes between age groups for most of the early responses. As an exception, the N21 response was enhanced, both in averaged waveform and in single-trial analysis, in the prepubertal rats compared to the adolescents.

The determination of peak latencies was based on fixed time windows, and it was thus not possible to examine the age-related changes in individual evoked response latencies at the single-trial level; this would have required adjusting the time windows individually. However, the overall complexity of auditory responses appeared earlier as age increased (see Figure 5). Therefore, the present findings do not allow us to conclude that there are no age-related changes in the timing of early auditory responses after 28 postnatal days in rodents. Further research is needed, with stimulation and analysis parameters optimized for early responses, to achieve a more accurate trajectory of latency changes by age.

A similar prolonged response in humans has already been reported in the 1970s (Picton et al. 1974), but in most of the earlier studies, the emphasis has been on cognitive processing or processing of different perceptual categories. Our late response was evoked in purely passive conditions without attention, priming, or oddball manipulation. Stronger or longer-lasting activation in this time window in purely passive conditions has been suggested to reflect a lower level of automatization or efficiency in cognitive tasks (Albrecht et al. 2000; Parviainen et al. 2011). However, other studies have linked stronger M250 responses with improved cognitive performance, for example, better phonological skills in the case of specific language impairment (van Bijnen et al. 2019) or higher response inhibition (van Bijnen et al. 2022). It is thus not straightforward to conclude whether increased activity in this time window should be considered behaviorally beneficial or an indicator of an immature and hence less efficient system. Understanding the developmental changes in the response properties of the cortical auditory pathway is critically needed before reaching a better understanding of the brain basis of higher-level neurocognitive development.

Based on our current findings, we propose that the unique prolonged response in the juvenile brain reflects increased excitability of cortical circuitry by external stimulation. The passively presented sine-wave tones used in the present study can be considered probes for examining the response properties of the cortical auditory pathway. Our results show that the synaptic response properties, which are generally considered to underlie the cortically measured evoked responses, significantly differ in the developing brain, still at the age of 13 in humans. Interestingly, when cortical reactivity has been probed, instead of natural sensory stimuli, by transcranial magnetic stimulation, the sensorimotor cortex exhibits similar prolonged reactivity to the M250 response in our study (Määttä et al. 2017).Together with these earlier findings for the sensorimotor cortex, as well as the evidenced prolonged activation pattern in the somatosensory cortex for infants (Pihko et al. 2009), our results imply that there may be a domain-general, and even species-general, delayed response type arising in the sensory cortices, which is characteristic of the maturing cortex.

What could be the underlying reason at the synaptic level for this prolonged activity? The recorded magnetic or electric field changes from outside the brain are directly associated with the local field potentials that arise from synaptic signaling in the laminar circuits. Thus, any developmental change in synaptic connectivity is likely to be reflected in the MEG/EEG fields, albeit with less spatial accuracy and only as a rough sum of the more detailed laminar-level events. However importantly, the temporal characteristics reflect the electric events associated with synaptic signaling, and it is the developmental changes in these postsynaptic currents that most likely give rise to the observed age-related differences.

Although excitatory feedforward connections are the main driveway for information transfer, protracted development of inhibitory circuits brings about precision and “computational power” to the circuitry, for example, by narrowing the synaptic integration window in the auditory cortex.^33^ The temporal properties of the inhibitory neurotransmitter receptors, which can be either fast ionotropic (GABA-A) or slower metabotropic (GABA-B) (Connors et al. 1988; MacDonald and Olsen 1994; Couve et al. 2000), provide a relevant source of electric activity considering the developmental changes in the cortically recorded responses. Using pharmacological manipulation and TMS, the evoked responses during the first 30 ms were suggested to reflect excitatory neurotransmission, whereas later peaks at 40–50 ms and >100 ms were associated with inhibitory neurotransmission either by GABA-A or GABA-B receptors, respectively (Ferreri et al. 2011; Premoli et al. 2014).

Temporal characteristics of GABAergic synapses are the same across cortical areas and presumably also across species (Bettler et al. 2004). Indeed, based on its temporal characteristics, the prolonged response in the juvenile auditory cortex could be related to developmental changes in GABA-B neurotransmission. In support of this interpretation, a few TMS-EEG studies in children demonstrate a later component at 200–300 ms, which differs both in amplitude and in spatial distribution from the adult response. This later child-specific activation was suggested to reflect developmental differences in GABA_B_ergic neurotransmission and/or in cortico-subcortical-cortical loops (Ferreri et al. 2017; Määttä et al. 2019). We thus propose that the late emerging activity in the auditory cortex in the present study reflects increased excitability of the synaptic circuitry in the juvenile state, possibly associated with modulations in GABAB-related neurotransmission. This characteristic may be fundamentally important for emerging cognitive skills and provide a signature of an increased state of plasticity in humans. However, further studies are needed to directly test the role of GABAergic signals in cortically measured evoked fields. The speed-up of the response latencies, especially in the early time window, could also reflect the influence of myelination, besides increased temporal precision of synaptic signaling along with the maturation of the GABA(A)ergic system.

The unique aspect of this study is the cross-species comparison. The two species were not recorded under exactly identical conditions; for humans, we used passive presentation of stimuli during the awake state, while the rats were anesthetized. However, passive response properties of the cortex are not assumed to be radically influenced by light anesthesia, and the age-group comparison within species should not be influenced by differences in measurement conditions. The cross-species comparison in the early time window is not straightforward due to the different timing of the individual peaks in the response between species and age groups. Moreover, although at a juvenile stage, the current study focused on relatively old subjects in both species, and the developmental changes in children under nine years or rodents under one month of age need to be explored in future studies.

To conclude, we provide evidence of the maturational steps of the late auditory activation in another mammalian species alongside data from humans, which confirms the hypothesis of the obligatory nature of the late activation in the developing brain. The findings at the level of single trials confirm that the enhanced and delayed activity recorded from outside the cortex both in juvenile humans and in rats reliably reflects response properties of the developing neural circuitry, rather than being a result of the averaging procedure. In fact, the late activation was not only evident at the single-trial level but also showed stronger trial-by-trial stability in human children than the early, adult-like 100-ms response. This implies that the late evoked activation does not reflect imprecise temporal accuracy in the circuitry but represents a separate prolonged synaptic event, which is likely to be meaningful and functional for the developing brain.

## ACKNOWLEDGMENTS

We are grateful to Joona Muotka for his technical support in the statistical analysis.

## SUPPLEMENTARY DATA

Appendix A. Average responses to sine-wave tones in rats.

Appendix B. The number of single trials in each age group in humans and rats. Appendix C. Single-trial time series in all individuals for humans and rats.

## AUTHOR CONTRIBUTIONS

Conceptualization, K.L., J.K., M.P., and T.P.; investigation, K.L., J.K., and T.P.; formal analysis, K.L. and L.P.; writing—original draft, K.L. and T.P.; writing—review & editing, J.K., L.P., R.U.M., and M.P.; methodology, L.P., R.U.M., and M.P.; software, R.U.M.; supervision, M.P. and T.P.; project administration and funding acquisition, T.P.

