## Supplemental_information for "Early stage of life is characterized by increased excitability of the auditory cortex in both humans and rats"

**A. Average responses to sine-wave tones in rats.** Time course of activation evoked by the sine-wave tones as recorded by the silver wire electrode from the surface of the auditory cortex in rats.

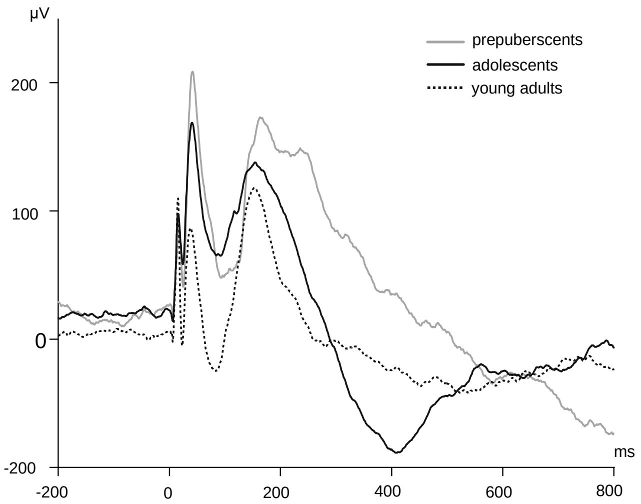


**B. The number of single trials in each age group in humans and rats.**

| | A<br>Mean<br>(Min–Max) | B<br>Mean<br>(Min–Max) | C<br>Mean<br>(Min–Max) |
|---|---|---|---|
| *Humans* | | | |
| M100 (N) | 69 (64–73) | 70 (61–73) | 75 (73–76) |
| M250 (N) | 74 (72–76) | 74 (70–76) | 74 (70–76) |
| *Rats* | | | |
| N6 (N) | 38 (30–44) | 37 (31–34) | 38 (31–41) |
| P14 (N) | 43 (26–54) | 42 (35–61) | 46 (39–61) |
| N21 (N) | 48 (34–62) | 41 (34–53) | 48 (39–55) |
| P41 (N) | 42 (29–51) | 47 (37–57) | 51 (40–64) |
| N108 (N) | 47 (37–56) | 46 (33–56) | 53 (44–63) |
| P214 (N) | 48 (34–56) | 52 (43–65) | 56 (43–66) |

**C. Single trial time series in all individuals for humans and rats.** Single-trial amplitude time courses of all individual subjects' auditory responses. The response amplitude values are illustrated with color codes across time in the horizontal direction, and consecutive trials are run in vertical columns starting from the first trial in the uppermost part of each subfigure. Units: individually scaled, relative to nanoamperometers in humans and relative to microvolts in rats. Plots are separated by species and by age group.

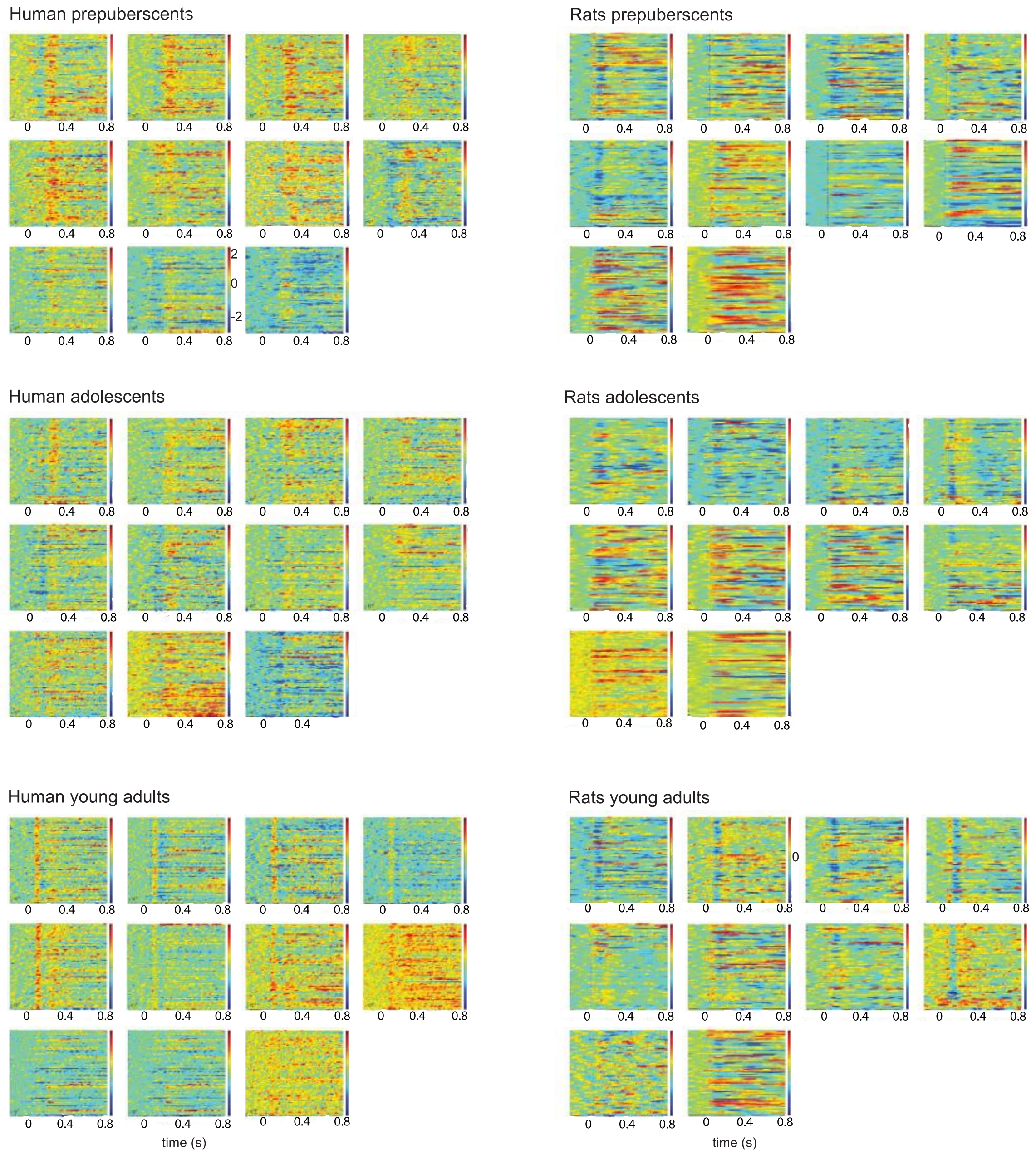
